# Auxin-inducible degron-mediated protein depletion in *Diplonema papillatum*

**DOI:** 10.64898/2026.09.11.751042

**Authors:** Gillian Clifford, Bungo Akiyoshi

## Abstract

Diplonemids are highly abundant unicellular flagellates that are evolutionarily divergent from traditional model eukaryotes. *Diplonema papillatum* is a model diplonemid for which several molecular tools have recently been established. However, no method for conditional protein depletion was available. Here, we have adapted an auxin-inducible degron (AID) system in *D. papillatum*. Depletion of the KIN-A motor protein (a glycomonad-specific component of the chromosomal passenger complex) causes defects in chromosome segregation and cell proliferation. The AID system also successfully depleted a divergent Mad1 protein that localizes at nuclear pores, and a flagellar component, CFAP20. Our study demonstrates the power of the AID system to perform loss-of-function studies in *D. papillatum*.

**Lay summary:** Diplonemids are single-celled microbes that live in the world’s oceans in enormous numbers, yet very little is known about their biology. They are only distantly related to humans and other well-studied model organisms used in cell biological research, but are close relatives of parasites that cause serious diseases such as African sleeping sickness, Chagas disease, and leishmaniasis. Studying diplonemids can therefore reveal both fundamental principles of cell biology and clues relevant to these related parasites. One diplonemid species, *Diplonema papillatum*, can be genetically manipulated in the lab, but researchers previously had no way to switch off individual proteins to study what they do. This work introduces such a tool, based on a plant-derived system that triggers controlled destruction of a target protein when a small chemical is added. We have used it to destroy three different proteins, showing that each is degraded efficiently and that one protein, called KIN-A, is required for cells to properly transmit their genetic material. This gives researchers a new way to study gene function in this understudied but important group of marine plankton.

## Introduction

Diplonemids are highly abundant but poorly understood marine organisms (1–3). They are evolutionarily very divergent from popular model eukaryotes that are widely used in cell biological research such as yeasts, worms, flies, and humans (4). Importantly, phylogenetic studies have established their close relationship with kinetoplastids, which include important pathogens (e.g. *Trypanosoma brucei, Trypanosoma cruzi*, and *Leishmania*) (5,6). In fact, diplonemids possess proteins that were initially thought to be specific to kinetoplastids, including a cytoskeletal protein, BILBO1 (7,8), some membrane-trafficking system components (9), and regulatory proteins involved in chromosome segregation (10,11). These findings highlight the close relationship between diplonemids and kinetoplastids (together called glycomonads (5)). Based on a number of unique biological features, Akiyoshi has proposed that glycomonads might be the earliest-branching eukaryotes (12,13), although this idea remains highly controversial (14). Regardless, the massive evolutionary distance between diplonemids and traditional model organisms means that studies of diplonemids could provide important insights into fundamental requirements of various biological processes in eukaryotes.

*Diplonema papillatum* (also known as *Paradiplonema papillatum* or *Isonema papillatum*) is a model diplonemid for which several molecular tools have recently been developed. A draft genome sequence is available (15). Genetic manipulation by means of homologous recombination enables epitope-tagging of a gene of interest at the endogenous locus (11,16,17). By contrast, there is no established method to perform conditional protein depletion, a critical tool for studying protein function, especially for essential genes whose knockout causes cell death (18). This represents a continuing drawback for functional studies in this organism.

The auxin-inducible degron (AID) system is a powerful tool to rapidly deplete specific proteins within cells (19). It takes advantage of a Skp1-Cullin-F-box (SCF) E3 ubiquitin ligase that contains the plant-specific F-box protein TIR1. Addition of auxin induces degradation of AID-tagged proteins by promoting an interaction between TIR1 and the degron, leading to polyubiquitination and rapid proteasomal degradation. In the AID2 system, expression of the TIR1 F74G mutant enables degradation of proteins fused to a 7-kDa degron tag (mAID) upon addition of the auxin analog 5-Ph-IAA (5-phenyl-indole-3-acetic acid) (20). The AID2 system was recently applied to *Trypanosoma brucei*, a kinetoplastid parasite that causes African trypanosomiasis, by expressing the entire SCF-TIR1 complex (21). Here, we establish the first conditional protein depletion method for *D. papillatum* by applying the AID2 system.

## Results

### Expression of OsTIR1 and OsSKP1 in *Diplonema papillatum*

To apply AID in *D. papillatum*, we engineered a cell line (DP183) that constitutively expresses TIR1 and SKP1 from rice, *Oryza sativa* (OsTIR1^F74G^ and OsSKP1). Like *T. brucei, D. papillatum* has polycistronic gene expression. We therefore designed a construct, encoding OsTIR1^F74G^ and OsSKP1, that integrates into an alpha-tubulin locus (Figure 1A). Immunoblotting confirmed expression of these proteins (Figure 1B). Treatment of these cells with 5 µM 5-Ph-IAA did not cause any growth defect (Figure 1C).

**Figure 1.**
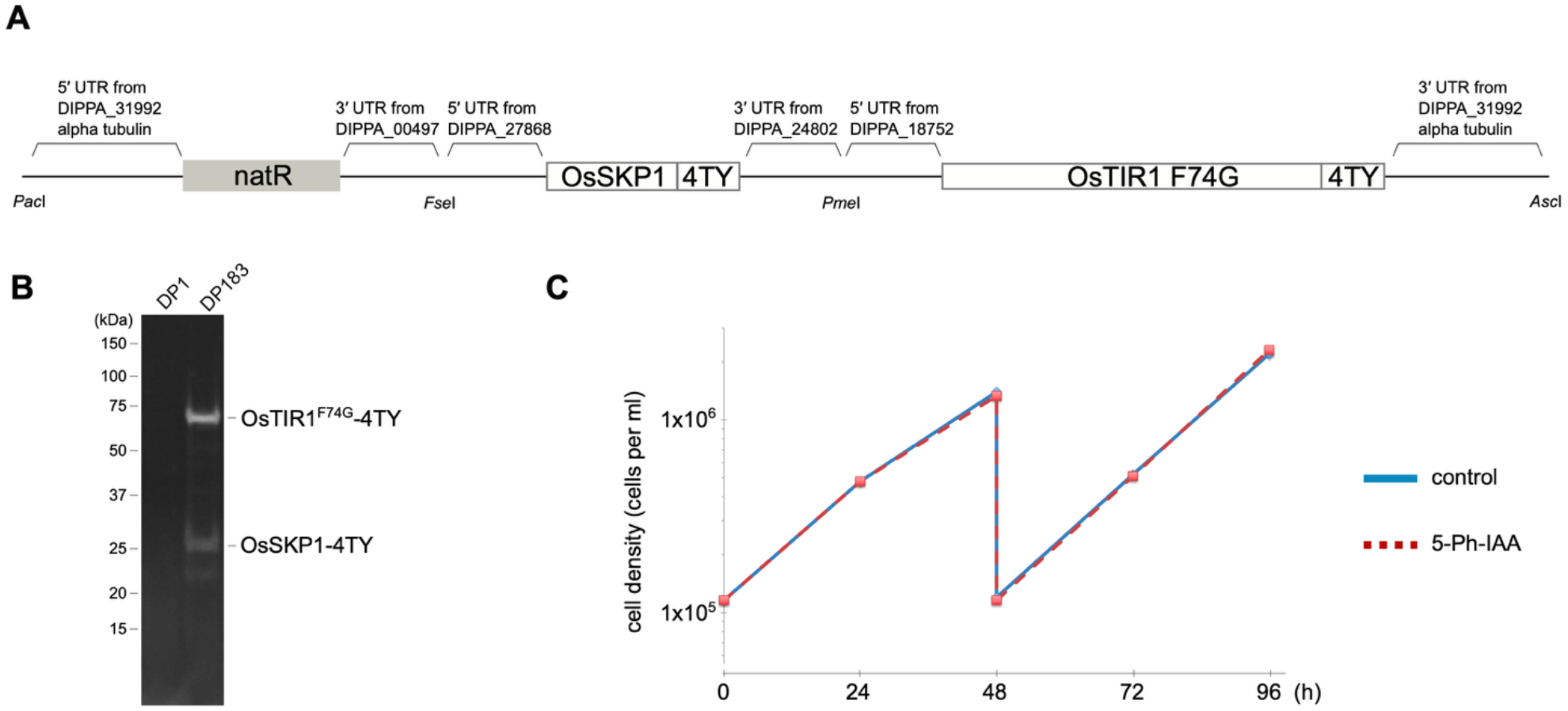
Expression of OsTIR1^F74G^ and OsSKP1 in *Diplonema papillatum*. (A) Schematic of the vector (pBA3611), which has a clonNAT selection marker and expresses OsTIR1^F74G^-4TY (theoretical molecular weight: 69 kDa) and OsSKP1-4TY (theoretical molecular weight: 24 kDa) from the alpha-tubulin locus (DIPPA_31992). (B) Immunoblotting with the BB2 anti-TY antibody shows expression of OsTIR1^F74G^ and OsSKP1 in DP183 cells, but not in the parental DP1 cell line. (C) Treatment with 5 µM 5-Ph-IAA does not cause any noticeable growth defect in DP183 cells. Control is untreated cells. The experiment was performed twice with similar results.

### Depletion of KIN-A fused with YFP-mAID causes growth defect

To examine whether AID works in *D. papillatum*, we chose KIN-A as a target because its homolog in *T. brucei* plays an essential role in chromosome segregation (10,22). To apply AID, we modified a YFP-tagging vector (11) to generate the YFP-mAID-tagging vector called pBA3622 (Figure 2A). Two homology arms were subsequently inserted into pBA3622 to facilitate the integration of YFP-mAID at the C-terminus of KIN-A at the endogenous locus in the DP183 cell line. Clonal cell lines expressing the fusion protein were screened by fluorescence microscopy, showing a nuclear localization pattern similar to KIN-A-YFP (11) (Figure 2B). After 4 h of treatment with 5 µM 5-Ph-IAA, KIN-A-YFP-mAID signal was markedly reduced (Figure 2B), and nuclear division defects were observed (Figure 2C). Importantly, 5-Ph-IAA treatment caused a severe growth defect in these cells (Figure 2D). Taken together, these results demonstrate that the AID2 system can efficiently deplete KIN-A-YFP-mAID and that KIN-A is essential for accurate chromosome segregation in *D. papillatum*.

**Figure 2.**
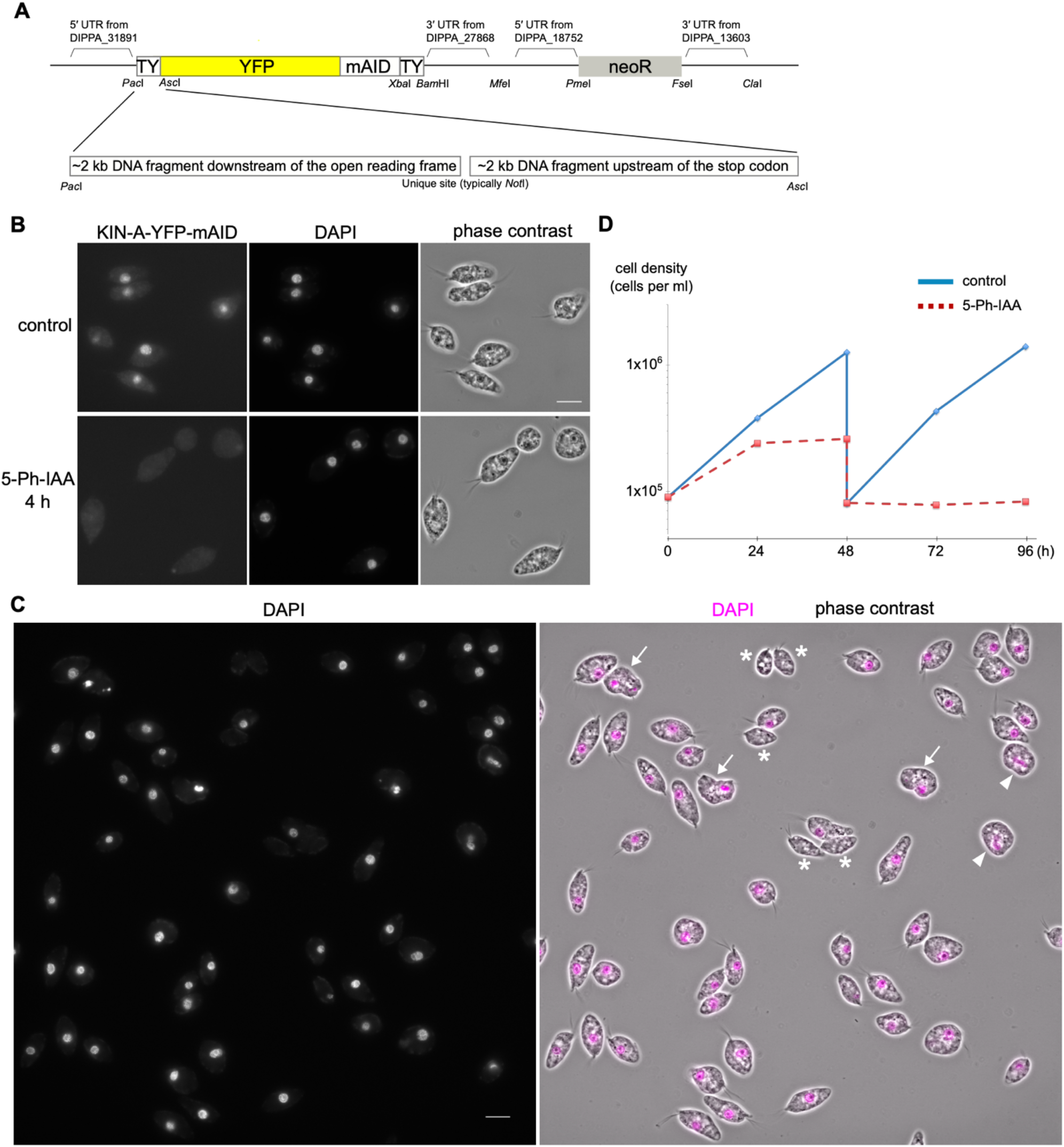
Depletion of KIN-A causes chromosome mis-segregation. (A) Schematic of the YFP-mAID tagging vector, pBA3622. For C-terminal tagging of a gene of interest, two homology arms are inserted into *Pac*I and *Asc*I sites of pBA3622. (B) Fluorescence microscopy shows reduction of KIN-A nuclear signal upon 5 µM 5-Ph-IAA treatment for 4 h. Control is untreated cells. (C) A wide field of view showing nuclear division defects upon 4-h treatment with 5-Ph-IAA. Asterisks indicate anucleate cells. Arrowheads and arrows indicate anaphase cells with lagging chromosomes or unequal DNA masses, respectively. (D) Growth curves of control and 5-Ph-IAA-treated cells show a severe growth defect after depletion of KIN-A. Cells were diluted on Day 2. The experiment was performed at least twice with similar results. Cell line, DP186. Scale bars: 10 µm.

### AID works for other nuclear and cytoplasmic proteins

We next tested two other targets: Mad1 and CFAP20. Mad1-YFP localizes at nuclear pores during interphase (11). The C-terminal YFP-mAID fusion for Mad1 showed a similar localization pattern (Figure 3A). CFAP20 (cilia and flagella associated protein 20, DIPPA_14323) is a highly conserved structural protein found in the axoneme (23). CFAP20-YFP-mAID localizes at flagella as expected (Figure 3B). Treatment with 5-Ph-IAA caused a reduction in YFP signal for both fusion proteins (Figure 3A,B). Severe growth defects were observed for Mad1 (Figure 3A) but not for CFAP20 (Figure 3B). These results demonstrate that the AID2 system enables conditional depletion of various targets in *D. papillatum*.

**Figure 3.**
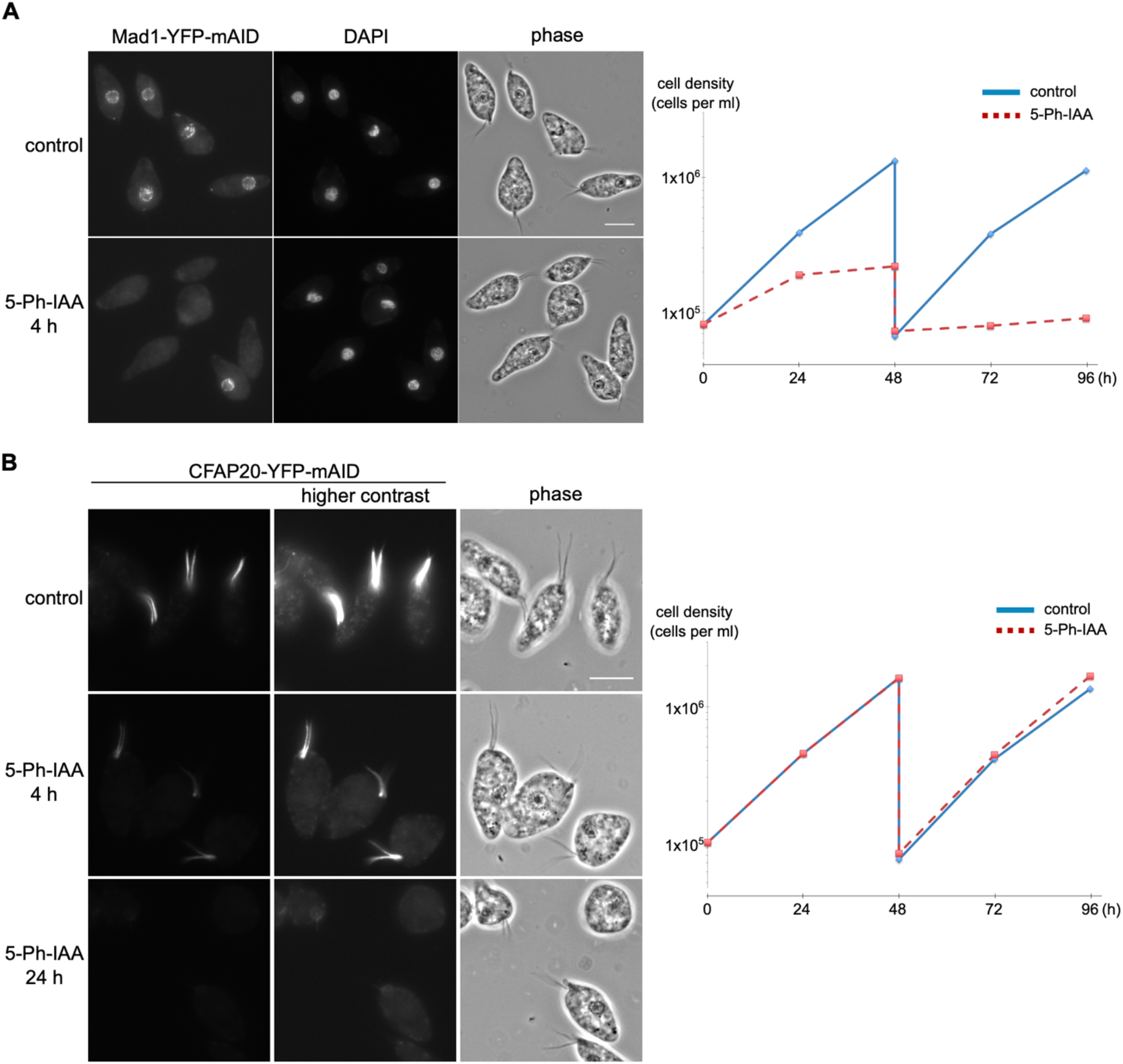
Depletion of Mad1 and CFAP20. Left: Fluorescence microscopy images of cells expressing Mad1-YFP-mAID (A) and CFAP20-YFP-mAID (B), showing reduction of YFP signals upon treatment with 5 µM 5-Ph-IAA. Right: Growth curves of control and 5-Ph-IAA-treated cells, showing that Mad1, but not CFAP20, is essential for proper cell growth. The experiment was performed twice with similar results. Cell lines, DP187, DP192. Scale bars: 10 µm.

## Discussion

Diplonemids are highly abundant and diverse flagellates that are widespread throughout the oceans of the world, especially in the deep sea (1,2). Although ∼10 diplonemid species have become culturable, *D. papillatum* remains the only genetically tractable diplonemid (3). Importantly, many proteins present in divergent diplonemids are conserved in *D. papillatum* (24), meaning that this model diplonemid could serve as an experimentally tractable proxy for the whole diplonemid group. The AID2 system described in this work allows researchers to perform conditional protein depletion to infer the function of a protein of interest in *D. papillatum*.

The doubling time of *D. papillatum* is 10–12 h at 27°C. For KIN-A, we observed a good signal reduction and chromosome segregation defects after 4 h of 5-Ph-IAA treatment, meaning that it is possible to examine the first mitotic event after KIN-A depletion. By contrast, it took longer to fully deplete CFAP20, likely due to limited accessibility of the SCF-TIR1 complex to flagella. Although no growth defect was observed for CFAP20-depleted cells, our preliminary observations indicate that they have motility defects, suggesting that flagellar motility is dispensable for cell growth, at least under laboratory growth conditions. It will be interesting to apply the AID2 system to other proteins in *D. papillatum* and observe their depletion kinetics and phenotypes.

In summary, the AID2 system establishes a general framework for functional characterization in *D. papillatum* and may be adaptable to other culturable diplonemid species as genetic tools become available for them.

## Materials and Methods

### Plasmids and primers

Plasmids used in this study are listed in Table 1. Sequences of primers, vectors, and synthetic DNA fragments are available in Figshare (https://doi.org/10.6084/m9.figshare.33545926). pBA3611 containing a synthetic DNA fragment (BAG276) encoding OsTIR1^F74G^-4TY, OsSKP1-4TY, and a selection marker, natR, was purchased from GeneArt. These three genes were codon optimized for expression in *Diplonema papillatum* (15). pBA3622 (TY-YFP-mAID-TY) was made in two steps. First, a synthetic DNA fragment encoding TY-YFP-AID-TY (BAG264) was cloned into pBA3294 using *Pac*I/*Bam*HI, creating pBA3575. Then a synthetic DNA fragment (BAG274) was subcloned into pBA3575 using *Asc*I/*Xba*I, creating pBA3622. The mAID gene was codon optimized for expression in *T. brucei* (25). C-terminal YFP-mAID tagging constructs were made using pBA3622 and a strategy similar to that used for C-terminal YFP tagging constructs using pBA3294 (11,26). Briefly, two ∼2 kb homology arms amplified from genomic DNA by PCR using KOD one polymerase (Merck) were inserted into pBA3622 cut with *Pac*I and *Asc*I, with a *Not*I site between the two PCR fragments. Plasmids were validated by Oxford Nanopore whole plasmid sequencing (Plasmidsaurus). As previously observed in our attempts to tag other genes (11), there were occasional mismatches between nanopore sequencing results and expected plasmid sequences, especially in repetitive sequences in 3′ UTR regions.

**Table 1.** Plasmids used in this study.

| Name | Description (Source) |
| --- | --- |
| pBA3294 | TY-YFP-TY tagging vector (11) |
| pBA3575 | TY-YFP-AID-TY tagging vector (this study) |
| pBA3611 | OsTIR1<sup>F74G</sup>-4TY, OsSKP1-4TY expression from an alpha-tubulin locus (this study) |
| pBA3622 | TY-YFP-mAID-TY tagging vector (this study) |
| pBA3642 | C-terminal YFP-mAID-TY tagging for KIN-A at the endogenous locus (this study) |
| pBA3694 | C-terminal YFP-mAID-TY tagging for Mad1 at the endogenous locus (this study) |
| pBA3699 | C-terminal YFP-mAID-TY tagging for CFAP20 at the endogenous locus (this study) |

### Diplonema cultures and C-terminal YFP-mAID-tagging of genes at the endogenous locus

All cell lines used in this study were derived from *Diplonema papillatum* (ATCC 50162) and are listed in Table 2. Cells were grown at 27°C in artificial sea water containing 36 g/L Instant Ocean Sea Salt (Instant Ocean), 1 g/L tryptone (Formedium, TRP01), and 1% fetal bovine serum (Merck, F9665) in vented flasks. Cell growth was monitored using a CASY cell counter (Roche).

**Table 2.** Cell lines used in this study.

| Name | Description (Source) |
| --- | --- |
| DP1 | Wild-type *Diplonema papillatum* (ATCC 50162) |
| DP183 | OsTIR1<sup>F74G</sup>-4TY, OsSKP1-4TY (this study) |
| DP186 | OsTIR1<sup>F74G</sup>-4TY, OsSKP1-4TY, KIN-A-YFP-mAID-TY (this study) |
| DP187 | OsTIR1<sup>F74G</sup>-4TY, OsSKP1-4TY, Mad1-YFP-mAID-TY (this study) |
| DP192 | OsTIR1<sup>F74G</sup>-4TY, OsSKP1-4TY, CFAP20-YFP-mAID-TY (this study) |

To obtain a clonal cell line, DP183, expressing OsTIR1^F74G^ and OsSKP1, 10 µg of pBA3611 was cut with *Pac*I and *Asc*I, cleaned up by ethanol precipitation, resuspended in transfection reagent (Ingenio Electroporation Kit for the EZporator Electroporation System, Cambridge Bioscience), and transfected into ∼2 × 10^7^ wild-type cells (ATCC 50162) using Amaxa Nucleofector IIb (Lonza Bioscience) as previously described (26). One day after electroporation, limiting-dilution isolation was performed with media containing 400 µg/mL clonNAT (2BScientific) to obtain clonal lines.

C-terminal YFP-mAID tagging constructs were transfected into DP183 as above using approximately 5 to 10 µg of plasmids linearized by *Not*I. Populations of transfected cells were obtained by addition of 75 µg/mL G418 (Merck). Once populations had recovered, then clonal lines were isolated by limiting dilutions and screened by microscopy for YFP-positive cells. 5-Ph-IAA (Cambridge Bioscience) was dissolved in DMSO to make a 5 mM stock solution and added to a final concentration of 5 µM in cell cultures.

### Immunoblot

Cells were harvested by centrifugation (2000 g for 3 min) and resuspended in 2x LDS sample buffer (Thermo Fisher Scientific) with 100 mM DTT. The samples were boiled for 3 min and loaded onto an SDS-PAGE gel (175 V for 30 min). The gel was transferred to a nitrocellulose membrane using a Trans-Blot Turbo Transfer System (Bio-Rad). The membrane was blocked with 3% milk in PBST for 30 min, followed by incubation with the anti-TY mouse monoclonal BB2 antibody (27) (1:100 dilution in PBST) at 4°C overnight with gentle shaking. The membrane was washed for 5 min three times in PBST, and incubated with the secondary antibody, IRDye 680RD goat anti-mouse (926-68070; LI-COR) (1:5000 dilution in PBST). After three washes, bands were visualized on a ChemiDoc Imaging System (Bio-Rad).

### Microscopy

To observe native YFP signals, cells fixed in formaldehyde were imaged as previously described with minor modifications (8). Briefly, 1 mL of cell culture was centrifuged at 2000 g for 3 min. Cells were fixed with 4% formaldehyde solution (Life Technologies, 28906) diluted in PBS for 5 min, rinsed with PBS once, resuspended in a small volume of DABCO mounting media (1% w/v 1,4-diazabicyclo[2.2.2]octane, 90% glycerol, 50 mM sodium phosphate pH 8.0) with 100 ng/mL DAPI, and mounted onto glass slides. Images were captured on an Axio Imager.Z2 microscope (Zeiss) equipped with ZEN software using a Hamamatsu ORCA-Flash4.0 camera with 63× objective lenses (1.40 NA). Twenty-five z-sections covering 6 µm were collected. Images were analyzed in ImageJ/Fiji (28). Figures were made in Inkscape (version 1.4, https://inkscape.org/).

## Data availability

### Underlying data

Figshare: Extended Data for “Auxin-inducible degron-mediated protein depletion in *Diplonema papillatum*” (https://doi.org/10.6084/m9.figshare.33545926)

This project contains the following underlying data:

- Table S1 (Excel file)
  ∘ Details of cell lines and how they were made
  ∘ Details of plasmids and how they were made
  ∘ Primer sequences
  ∘ DNA sequences for pBA3294, pBA3575, pBA3611, pBA3622, BAG264, BAG274, BAG276
- Table S2 (Excel file): Raw data for growth curves
- DP1_DP183_Immunoblot.tif: Raw data for immunoblot
- Raw microscopy images (TIFF files)
  ∘ DP186_control.tif: KIN-A-YFP-mAID, untreated control
  ∘ DP186_4h.tif: KIN-A-YFP-mAID, 4-h treatment with 5-Ph-IAA
  ∘ DP186_4h_wide_field.tif: KIN-A-YFP-mAID, 4-h treatment with 5-Ph-IAA
  ∘ DP187_control.tif: Mad1-YFP-mAID, untreated control
  ∘ DP187_4h.tif: Mad1-YFP-mAID, 4-h treatment with 5-Ph-IAA
  ∘ DP192_control.tif: CFAP20-YFP-mAID, untreated control
  ∘ DP192_4h.tif: CFAP20-YFP-mAID, 4-h treatment with 5-Ph-IAA
  ∘ DP192_24h.tif: CFAP20-YFP-mAID, 24-h treatment with 5-Ph-IAA

## Acknowledgments

We thank Keith Gull for the BB2 antibody. Bungo Akiyoshi was supported by a Wellcome Discovery Award (227243/Z/23/Z).

## Author contributions

G.C. performed DNA cloning. B.A. made transgenic lines, performed imaging, and wrote the manuscript.

## Rights retention

This research was funded in whole, or in part, by the Wellcome Trust (227243/Z/23/Z). For the purpose of open access, the author has applied a CC BY public copyright licence to any Author Accepted Manuscript version arising from this submission.

